# Neph/Nephrin-like adhesion and tissue level pulling forces regulate cell intercalation during *Drosophila* retina development

**DOI:** 10.1101/564708

**Authors:** Laura Blackie, Melda Tozluoglu, Mateusz Trylinski, Rhian F. Walther, Yanlan Mao, François Schweisguth, Franck Pichaud

## Abstract

Intercalation between neighboring cells contributes to shaping epithelial tissues and is regulated by the contractile actomyosin cytoskeleton. While intercalation typically occurs over minutes, instances of much slower cell intercalation have been reported during organogenesis. This is observed, for example, for the four glial-like cone cells (CC) that intercalate during *Drosophila* retinal patterning. Here we show that Myosin-II activity in the CCs is largely dispensable for their intercalation. Instead, we find that differential activity of the Notch-signaling pathway within the CC quartet regulates intercalation, which also depends on the cell adhesion proteins Roughest and Hibris. In addition, mathematical modeling predicts that forces external to the intercalating CC quartet are necessary for intercalation. Consistent with this prediction we show that the surrounding primary pigment cells are under significant contractile tension. Altogether, our work elucidates a novel mode of cell intercalation that relies on Neph/Nephrin-like adhesion and forces external to the intercalating cells.

## INTRODUCTION

Epithelial tissues line most of our organs and consist of epithelial cells that are polarized along the apical (top) basal (bottom) axis and are attached to each other through lateral cell-cell junctions called *Adherens Junctions* (*AJ*). Next to mediating intercellular adhesion, *AJs* also support mechanical coupling between cells (Lecuit and Yap, 2015). During development, steps of neighbor exchange through polarized cell intercalation can promote tissue elongation, as seen in the *Drosophila* germband (Bertet et al., 2004; Blankenship et al., 2006; Tetley et al., 2016; Zallen and Wieschaus, 2004), convergent extension during mouse gastrulation (Yin et al., 2008) and renal tube development in mice (Karner et al., 2009) or *Xenopus* (Lienkamp et al., 2012). In addition, MyoII promotes the intercalation of newly specified photoreceptors to assemble the ommatidium, which is the physiological unit of the retina (Robertson et al., 2012). In all these contexts, most steps of cell intercalation that have been studied occur over tens of minutes.

The mechanisms of germband elongation in *Drosophila* are particularly well understood. In this tissue, intercalation occurs between four cells or higher order six-cell multicellular rosettes. The orientation of intercalation is regulated by upstream spatial information, which instructs dorso/ventral (D/V) *AJs* to shrink and subsequent antero/posterior (A/P) *AJs* to elongate (Pare et al., 2014; Zallen and Wieschaus, 2004). A polarized cytosolic contractile actomyosin meshwork present in the A and P cells pulls on the D/V junctions (Levayer and Lecuit, 2013; Rauzi et al., 2010) to power shrinkage of the D/V AJs. Concomitantly, non-muscle Myosin-II (Myo-II) accumulates at the D/V *AJ*s where it ratchets the contractile forces applied onto these *AJ* by the cytosolic contractile actomyosin meshwork (Bertet et al., 2004; Blankenship et al., 2006; Rauzi et al., 2010; Zallen and Wieschaus, 2004). In the germband, the exact relationship between the pulsatile-contractile actomyosin meshwork and the *AJ* pool of actomyosin is not fully understood, but several key factors are involved. Typically, the RhoA-Rok pathway acts upstream of Myo-II planar polarization and activity (Kasza et al., 2014; Munjal et al., 2015; Simoes Sde et al., 2010; Simoes Sde et al., 2014). Notably, polarization of the Guanine nucleotide exchange factor RhoGEF2 plays an important role as it activates RhoA and the Rok-MyoII pathway at the shrinking D/V *AJ*, which is depleted for the polarity protein Bazooka (Baz/Par3) (Levayer et al., 2011; Simoes Sde et al., 2010; Simoes Sde et al., 2014).

Epithelial cell intercalation can also be less deterministic than that documented in the germband. In the fly notum for example, intercalation is stochastic and reversible, and seems to be required to promote tissue fluidity (Curran et al., 2017). While in this tissue, *AJ* remodeling depends on Myo-II, increased accumulation of Myo-II at the cell *AJs* acts as part of a brake to stop intercalation as the tissue reaches its final configuration. Similarly, in the dorsal branch of the fly trachea where cells intercalate to form tubes, Myo-II is largely dispensable and instead, forces promoting intercalation between branch-cells are generated by the migrating tip-cell, which pulls on the branch cells (Ochoa-Espinosa et al., 2017). In these two model systems, a role for the cytosolic contractile actomyosin machinery has not been investigated.

While in simple epithelia, cell intercalation is an essential mechanism of tissue morphogenesis, we know much less about the mechanisms that regulate the morphogenesis of organs, which typically consist of multiple cell types. The fly retina and its basic physiological sensory unit, called the ommatidium, is an excellent model system to study the mechanisms of organogenesis. It is a highly stereotyped, multicellular hexagonal structure that consists of four core glial-like cells (cone cells, CC), surrounded by two epithelial-like primary pigment cells (PPC), themselves surrounded by a complement of six secondary and three tertiary epithelial-like pigment cells (collectively referred to as Inter-Ommatidial Cells; IOCs), and three mechano-sensory bristle organs (Cagan, 2009; Cagan and Ready, 1989; Charlton-Perkins et al., 2017; Ready et al., 1976; Wolff., 1993) (Figure 1A). Altogether, approximately 750 ommatidia form the compound eye. While they assemble into a cell quartet atop the photoreceptors, the CCs also recruit the two PPCs to the ommatidium through cell-cell interactions mediated by Notch (N). The anterior (A) and posterior (P) CC express the Notch-ligand Delta (Dl), which is thought to signal to neighbors to differentiate as PPCs (Nagaraj and Banerjee, 2007). Shortly after PPC recruitment, the CCs then undergo intercalation such that the polar (Pl) and equatorial (Eq) CCs come into contact as they intercalate between the A/P CCs (Figure 1B). However, a main departure point from instances of intercalation that have been previously studied is that CC intercalation is slow and occurs over hours. Mechanisms of actomyosin contraction typically operate over minutes, suggesting CC intercalation might rely on alternative mechanisms. Alternative mechanisms might include cell sorting or adhesion. For instance, patterning of the ommatidium requires preferential adhesion between the PPCs and surrounding IOCs. The PPCs express the Nephrin-like immunoglobulin adhesion protein Hibris (Hbs) that is thought to bind the homologue of Neph-1, Roughest (Rst), which is expressed in the IOCs (Bao and Cagan, 2005). Furthermore, Hbs regulates CC intercalation (Grillo-Hill and Wolff, 2009).

**Figure 1:**
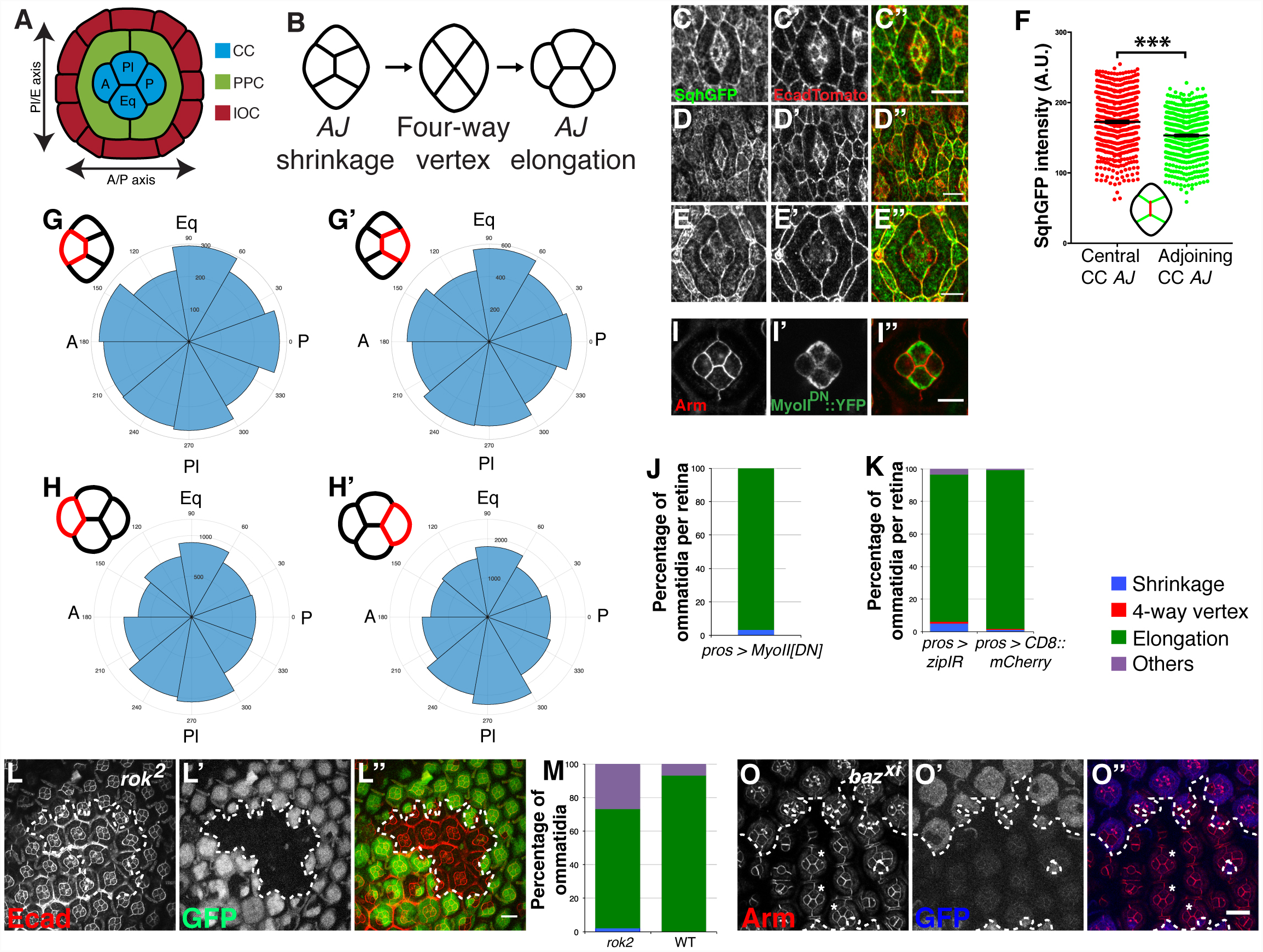
Myosin-II activity in the CC is not essential for their intercalation. **(A)** The arrangement of cells in the ommatidium. **(B)** Stages of intercalation of the glial-like CCs**. (C**-**E)** *sqh*^*AX3*^*;sqh-sqh::GFP/+;Ecad::Tomato/+* flies showing localization of Myo-II (Sqh::GFP) at the junction shrinkage **(C)**, four-way vertex **(D)** and junction elongation **(E)** stages of CC intercalation. **(F)** Quantification at the *AJ* shrinkage stage of Sqh::GFP intensity on shrinking A/P-CC *AJ* compared to adjoining CC-CC *AJs* paired by ommatidium (n=514 ommatidia). Paired T-test: p<0.0001. **(G-H)** Polar histograms showing directions of Myo-II flow vectors calculated by PIV. **(G)** A-CC and **(G’)** P-CC during A/P-CC *AJ* shrinkage. **(H)** A-CC and **(H’)** P-CC during Pl/Eq-CC *AJ* elongation. **(I)** *UAS-MyoII*^*DN*^::YFP expressed under control of *pros-Gal4*. **(J)** Progression of CC intercalation at 29°C when MyoII^DN^ is expressed in the CCs (n=4 retinae, 2212 ommatidia). **(K)** Progression of CC intercalation for *UAS-zipIR* expressed under control of *pros-Gal4* alongside matched controls expressing *UAS-CD8::mCherry*, raised at 29°C. (*pros-Gal4; zipIR*: n=6 retinae, controls: n=4 retinae).**(L)** *rok*^*^2^*^ mutant clones marked by the lack of GFP in a wild-type background. **(M)** Progression of CC intercalation at 40%APF within clones (*rok*^*^2^*^ mutant, n=138 ommatidia) compared to surrounding wild-type tissue (n=333 ommatidia). **(O)** *baz*^*xi*^ mutant tissue marked by the absence of GFP. Mutant ommatidia comprised of 4 glial-like CCs complete intercalation (indicated with a white star). Scale bars: **(C-E,I)** = 5μm, **(L,O)** = 10μm. Error bars = S.E.M.

Here we combined molecular genetics, laser-mediated cell contact ablation and *in silico* modeling to elucidate the mechanisms of slow CC intercalation during ommatidium morphogenesis. Altogether, our work reveals a new function for the N-signaling pathway in regulating adhesion-based cell intercalation through the Hbs/Rst system. Our mathematical modeling, which is verified by experimental data, also shows that CC intercalation requires contractile forces outside of the intercalating CC quartet. This novel mode of cell intercalation does not rely on MyoII activity in the intercalating cells. Instead it requires the combination of Neph/Nephrin-like adhesion intrinsic to the intercalating cells and between these cells and the surrounding PPCs. Our computer simulation of CC intercalation using a vertex model of the developing ommatidium also predicts a requirement for pulling forces outside of the intercalating cell quartet. Consistent with this predicted requirement, we show that the surrounding PPCs are highly contractile.

## RESULTS

### Myosin-II activity in the CC is not essential for their intercalation

CC intercalation unfolds over 10hrs (Movie S1), which is more than ten times longer than it takes for a group of four cells to intercalate in the germband. To ask whether MyoII regulates CC intercalation, we analyzed the intensity of Myo-II at the *AJs* (Figure 1C-E). Examining the CC quartet, we could detect only a marginal Myo-II enrichment at the shrinking A/P-CC *AJ*, when compared to the other CC *AJs* (Figure 1F). In addition, no Myo-II flows could be detected that were directed toward the shrinking A/P-CC *AJ* (Figure 1G), nor later on, with respect to the elongating Pl/Eq-CC *AJ* (Figure 1H).

Next, we sought to perturb Myo-II in the CCs using a CC-specific Gal4 driver (*pros-Gal4*) to express a dominant negative version of the Myo-II heavy chain, Zipper^DN^::YFP (Barros et al., 2003). This perturbation had very little effect on CC intercalation (Figure 1I-J). To complement these experiments, we made use of RNAi (IR) against Zipper (*zip*). Expression of *zipIR* did not significantly interfere with intercalation when expressed specifically in the CCs (Figure 1K). Ruling out an essential role for the Rok-Myo-II pathway and Baz during CC intercalation, we found that neither loss of *rok* (Figure 1L-M) nor *baz* function affected CC intercalation (Figure 1O). In the case of *baz*^*xi*^ loss-of-function, many ommatidia presented missing CCs or super-numerous CCs, which prevented us from scoring intercalation. However, all ommatidia containing a CC quartet had undergone intercalation. Together with our Myo-II measurements, these experiments indicate that Rok, Myo-II and Baz are largely dispensable for CC intercalation.

### CC intercalation occurs perpendicular to the elongation axis

In order to characterize the dynamics of CC intercalation, we used E-cad::GFP to visualize their perimeters in live preparations. As the CCs intercalate, ommatidial cells refine their shapes (Movie S1 and Figure 2A) and the average apical area of the CCs increases (Figure 2B). Quantification revealed that concurrent with A/P-CC *AJ* shrinkage, the CC cluster elongates along the Pl/Eq axis of the ommatidium, even though CC-CC *AJ* shrinkage takes place along this axis (Figure 2D-E). Furthermore, during the subsequent phase of intercalation, as the Pl/Eq-CC *AJ* elongates, we found that the CC quartet widens along the axis of *AJ* elongation (A/P axis of the ommatidium). Considering morphogenetic events outside of the CC quartet, we note that during the timeframe of CC intercalation, the *AJs* connecting the two PPCs lengthen at a higher rate than the CC-CC *AJs* (Figure 2C). This lengthening negatively correlates with the decrease in A/P-CC *AJ* length (Supplementary Figure 1A). On the other hand, we found that fluctuations in PPC-PPC *AJ* length do not correlate with fluctuations in the length of the shrinking A/P-CC *AJ* (Supplementary Figure 1B). Therefore, no direct coupling between these two sets of *AJs* seems to be taking place. However, quantifications revealed a cross-correlation between the shrinkage of the A/P-CC *AJ* and the expansion of the remaining adjoining CC-CC *AJs* (Figure 2F) but not between that of the CC-PPC *AJs* (Figure 2G). These results suggest that a mechanism operates whereby loss of *AJ* material at the shrinking A/P-CC contact is accompanied by apposition of *AJ* material at the adjoining CC-CC *AJs*. One possibility is that endocytosis and recycling of *AJ* material regulates this concomitant *AJ* shrinkage and lengthening.

**Figure 2:**
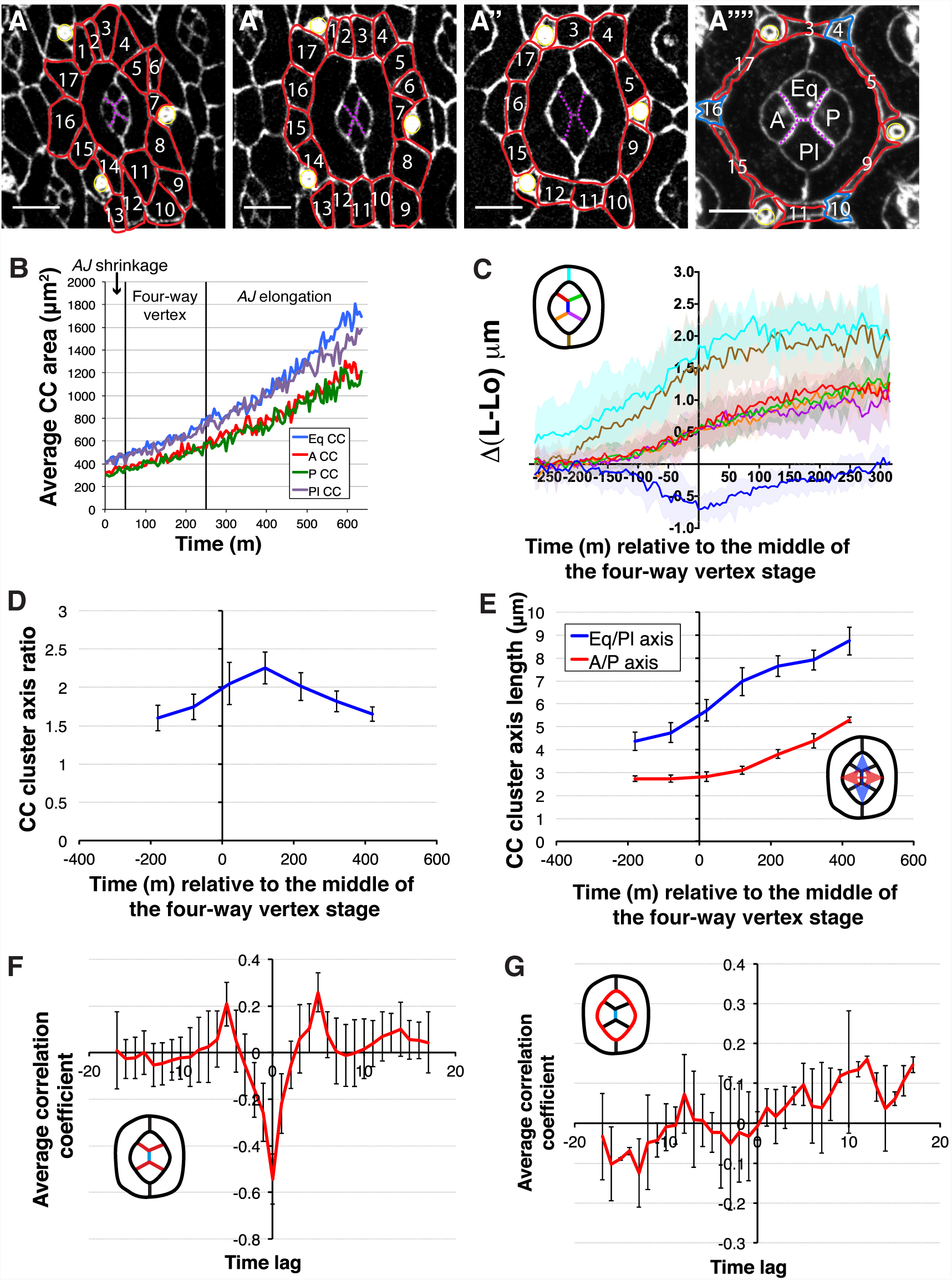
CC intercalation occurs perpendicular to the elongation axis. **(A)** Confocal sections taken from a time-lapse movie of ommatidium development, with *AJs* labeled with endogenous Ecad::GFP. IOCs are outlined in red and numbered through subsequent frames. Tertiary IOCs are labeled in blue. **(B)** Average apical area of CCs over time (n=4 ommatidia). Vertical lines demarcate the stages of CC intercalation. **(C)** Average relative length (L-L0) of CC and PPC *AJs* during ommatidium development. Time 0 is middle of the four-way vertex stage (n=13 ommatidia from 2 retinae). **(D)** Average CC cluster axis ratio over time relative to the middle of the four-way vertex stage (n=14 ommatidia). **(E)** Average CC cluster axes lengths over time relative to the middle of the four-way vertex stage (n=14 ommatidia). **(F)** Average cross-correlation of rate of change in the length of the central CC-CC *AJ* (shown in blue in the schematic), with the adjoining CC-CC *AJ* (shown in red in the schematic). Correlation coefficient: r=-0.54+/-0.11 (mean+/-S.D.) at a time lag of 0 (n=13 ommatidia). **(G)** Average cross-correlation of rate of change in length of the central CC-CC *AJ* (shown in blue in the schematic) with the CC-PPC *AJ* (shown in red in the schematic) (n=13 ommatidia). Scale bars = 5μm. Error bars = S.D.

### CC intercalation requires endocytosis

To test the requirement for endocytosis we made use of the thermo-sensitive Shibire^ts1^ protein (ortholog of *dynamin*). Inhibition of endocytosis specifically in the CCs from 20-40%APF caused the CC *AJs* to become convoluted (Figure 3A). This was specific to the *AJs* shared between the CCs, as the CC-PPC *AJs* were not affected. In addition, we observed an increase in the relative area occupied by the CC quartet. Interestingly, inhibition of endocytosis from 24-40% APF caused a stalling of CC intercalation at the early *AJ* shrinkage stage in 72% of the scored ommatidia, and at the four-way vertex in 18% of the scored ommatidia (Figure 3B-C). In order to refine this analysis, we transiently inhibited endocytosis for 4hrs during each of the three stages of CC intercalation: 20%APF (A/P-CC *AJ* shrinkage), 24%APF (four-way vertex) and 28%APF (Pl/Eq-CC *AJ* elongation (Figure 3D-F). In all three cases, CC intercalation was stalled at either the A/P-CC *AJ* shrinkage and/or four-way vertex stage (Figure 3G). Endocytosis inhibition at 24%APF had the greatest impact on CC intercalation, whereas inhibition at 20%APF and 28%APF had milder comparable effects (Figure 3G). Altogether these results demonstrate that endocytosis is required in the CCs in order to shrink the A/P-CC *AJ*.

**Figure 3:**
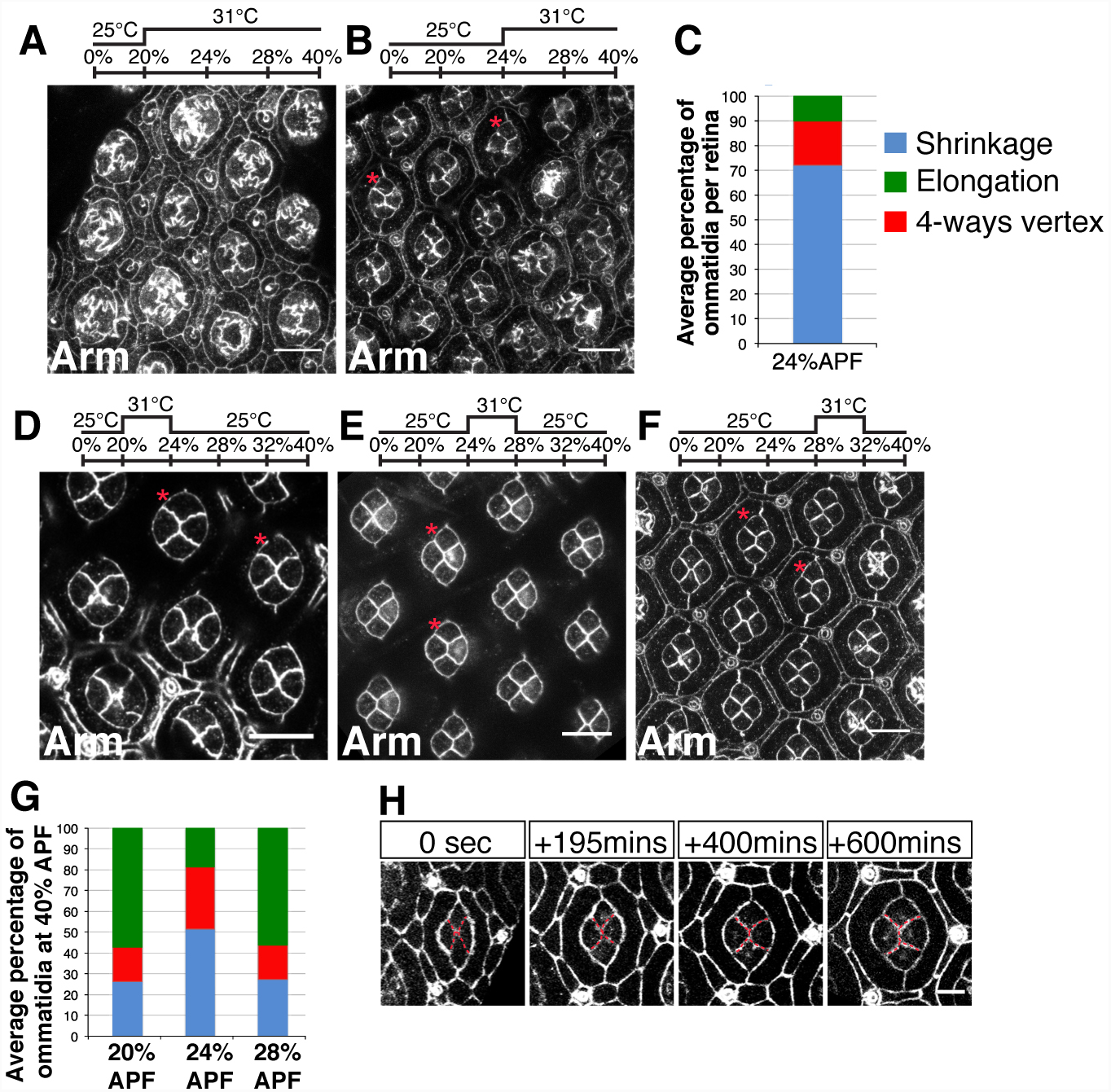
CC intercalation requires endocytosis. **(A-B,D-F)** Retina expressing *UAS-shibire*^*ts*^ under the control of *prosGal4* stained for Arm. Flies were raised at 25°C and then transferred to 31°C at **(A)** 20%APF, **(B)** 24%APF and incubated O/N. Flies were transiently transferred to restrictive temperature for 4hrs at **(D)** 20%APF, **(E)** 24%APF and **(F)** 28%APF. **(C)** Progression of CC intercalation in **(B)** (n=4 retinae, 1442 ommatidia). **(G)** Progression of CC intercalation in **(D-F)** (n=7, 6, 6 retinae respectively; 3352, 3100, 2797 ommatidia respectively). **(H)** Stills taken from a movie of retina expressing *UAS-shibire*^*ts*^ under the control of *prosGal4* with *Ecad::GFP* to label the *AJs.* CC *AJs* are overlaid in red. Scale bars: **(A-B,D-F)** = 10μm, **(H)** = 5μm.

Inhibition of endocytosis in the CCs could cause a stalling of cell intercalation or affect the stability of intercalation, causing intercalation to reverse. In order to distinguish between these possibilities, we performed time-lapse imaging of flies expressing *UAS-shibire*^*ts*1^ in the CCs. We found that inhibition of endocytosis caused a failure in A/P-CC *AJ* shrinkage (Movie S2). However, we also noted that CCs at the four-way vertex stage could revert to the initial CC configuration (Figure 3H). Therefore, endocytosis is required at multiple steps of CC intercalation.

### Roughest and Hibris are required for CC intercalation

Next to mechanisms of Myo-II contractility, adhesion is an essential regulator of cell shape (Hayashi and Carthew, 2004). Preferential adhesion through Rst-Hbs regulates the intercalation of the IOC during ommatidial patterning (Bao et al., 2010). We hypothesized that heterophilic adhesion through Hbs and Rst might be a good candidate mechanism for regulating CC intercalation. To test this hypothesis, firstly, we made use of endogenously tagged Rst::GFP and Hbs::GFP to assess their expression in CCs (Figure 4A-F). We found that both these adhesion molecules were localized at the CC *AJs* shared with the surrounding PPCs. In addition, we noted that both Rst and Hbs were also detected at the *AJ* shared by the two PPCs (Figure 4A-F). Secondly, to directly assess the requirement for Hbs and Rst during CC intercalation, we decreased their expression using RNAi lines in mosaic ommatidia (Figure 4G-K). These mosaic analyses confirmed that Hbs is required in all CCs for them to achieve their final configuration (Grillo-Hill and Wolff, 2009) (Figure 4G-G’’’ quantified in 4J). We obtained a range of phenotypes, including failure to shrink the A/P-CC *AJ* (Figure 4G’’), intercalation stalled at the 4-way vertex stage (Figure 4G’), and more profound defects in CC configuration suggesting CC sorting defects (Figure 4G’’’). Notably, decreasing the expression of *hbs* in the Eq-CC and Pl-CC led to limited defects in intercalation, while decreasing its expression in the A- or P-CC led to the stronger defects in CC configuration reminiscent of cell sorting defects (Figure 4G’’’ and 4J). These sorting defects consisted of the A- or P-CC minimizing their interface with their PPC neighbor, indicating in the A- and P-CC, Hbs is part of a preferential adhesion system with the surrounding PPCs. Therefore, during CC intercalation, there is a differential requirement for Hbs in the Pl/Eq-CCs and A/P-CCs. Next, we examined the requirement for Rst and found that decreasing its expression in the Pl- and Eq-CCs interfered with CC intercalation in 10% of the cases (Figure 4H-H’, quantified in 4K). These experiments suggest that an adhesion code, which includes Rst and Hbs is at play during CC intercalation and our mosaic analyses suggest that part of the instructive code resides in Rst being specifically required in the Pl-and Eq-CCs.

**Figure 4:**
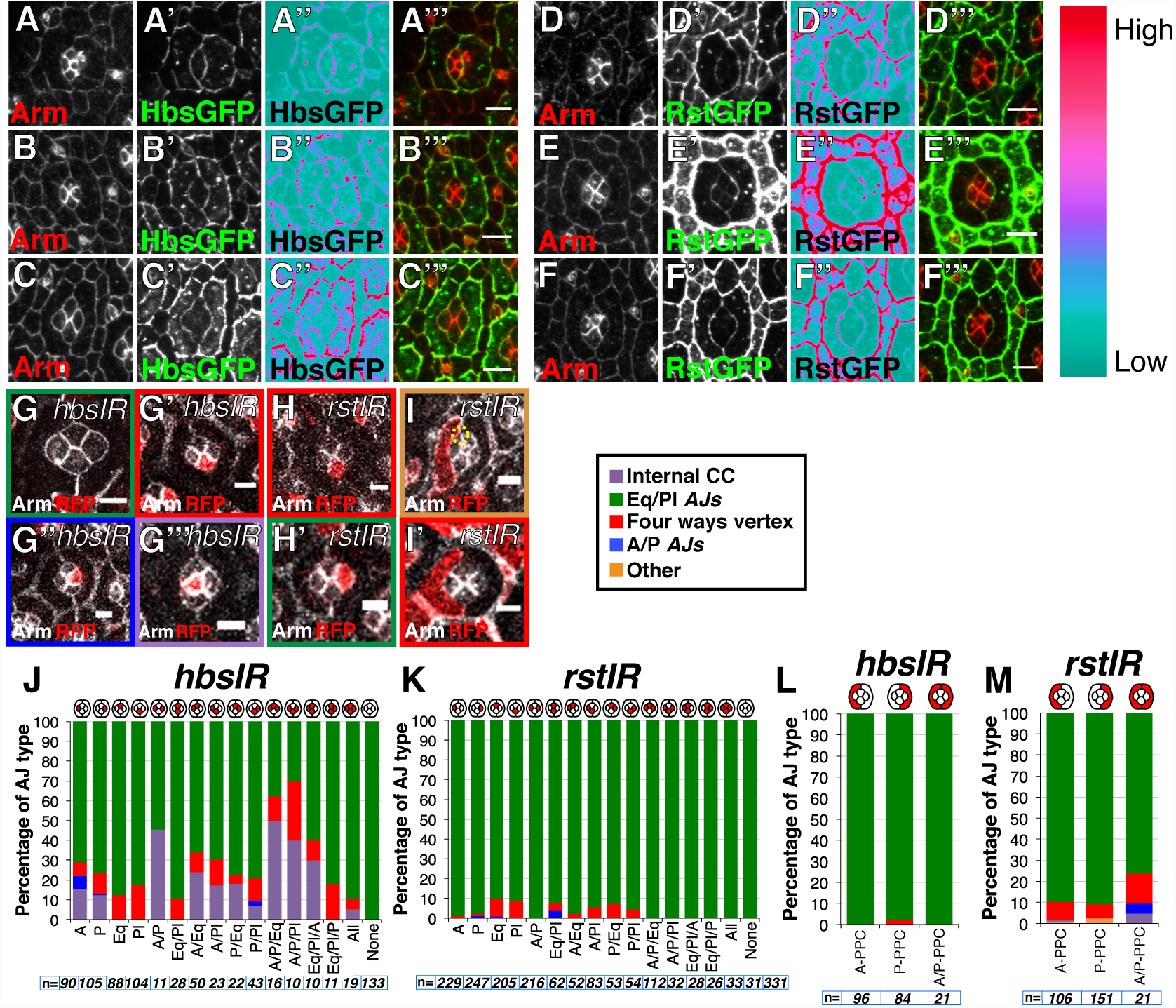
Roughest and Hibris are required for CC intercalation. **(A-F)** Confocal projection through the CCs showing **(A-C)** Hbs::GFP and **(D-F)** Rst::GFP at **(A,D)** *AJ* shrinking stage, **(B,E)** four-way vertex stage and **(C,F)** *AJ* elongation stage. **(A”,B”,C”,D”,E”,F”)** LUT to visualize variation in levels along the CC-PPC *AJ*. Note the reduction in intensity around the Pl and Eq CCs. **(G)** Representative wild-type, control ommatidium from a *hbsIR* mosaic retina **(G’)** Eq cell expressing *hbsIR* (red) and stalled at the four-way vertex **(G’’)** P-CC expressing *hbsIR* and stalled at the A/P-CC *AJ* shrinking stage **(G’’’)** A-CC expressing *hbsIR* and showing a cell sorting phenotype. **(H)** Eq-CC expressing *rstIR* and stalled at the four-way stage **(H’)** P-CC expressing *rstIR* undergoing normal intercalation. **(I)** A-PPC expressing *rstIR* showing a displacement of the PPC-PPC *AJ* toward the vertex between the A/Pl-CC and the PPCs (circled with a yellow dashed line). **(I’)** A-PPC expressing *rstIR* and showing CC intercalation stalled at the four-way vertex. **(J-K)** Quantification of the percentage of ommatidia with each *AJ* type (A/P, four-way vertex, Pl/Eq, internal CC) when different combinations of CCs express **(J)** *hbsIR* and **(K)** *rstIR*. **(L-M)** Quantification of the percentage of ommatidia with each *AJ* type (A/P, four-way vertex, Eq/Pl, internal CC), when different combinations of PPCs express **(L)** *hbsIR* and **(M)** *rstIR*. N numbers are shown with each panel. Scale bars = 5μm.

### CC intercalation is regulated by the neighboring PPCs

Most Rst-Hbs staining in the CC is detected at their *AJs* with the surrounding PPCs (Figure 4C’’-4F’’). This suggests that the PPCs might influence CC intercalation through these proteins. To test this suggestion, we examined ommatidia presenting one or both PPCs deficient for *hbs* or *rst*. Our mosaic experiments using RNAi confirmed that Hbs is not required in the PPCs to promote CC intercalation (Grillo-Hill and Wolff, 2009) (Figure 4L). However, we found that when Rst expression is decreased in one PPC (A or P), or both the A/P-PPCs, CC intercalation was affected (Figure 4I-I’, quantified in 4M). A role for Rst in the PPCs is consistent with these cells presenting Rst at their *AJ* with the CC, where it would bind Hbs in *trans*. Altogether, these results suggest that the Hbs-Rst interaction at the CC/PPC *AJ* regulates the intercalation of the CC. We also note that in some cases, decreasing the expression of *rst* in a PPC leads to a shift in the position of the PPC-PPC *AJ* with respect to the Pl/Eq axis of the ommatidium (Figure 4I). This is consistent with Rst in the PPC regulating adhesion between the PPC and Pl/Eq-CC.

### Notch signaling regulates CC intercalation

Previous studies have shown that the expression of Hbs in the PPCs is regulated by Notch (N) (Bao, 2014). This link between the Neph/Nephrin-like adhesion system and N-signaling prompted us to investigate the role of N during CC intercalation. Firstly, we examined the distribution patterns of N and its ligand Dl before and after CC intercalation using live imaging of functional GFP-tagged proteins (Figure 5A-B) (Corson et al., 2017; Trylinski et al., 2017). At the onset of CC intercalation, we found that N was present at the apical cortex of all CCs (Figure 5A) whereas Dl was highly expressed in the A-CC and, albeit to a lesser extent, in the P-CC (Figure 5B). Dl levels were minimal in the Pl- and Eq-CC. We also examined the distribution of Neuralized (Neur), a developmentally-regulated E3 ubiquitin ligase that is required for Dl endocytosis and N receptor activation (Figure 5C) (Perez-Mockus et al., 2017a; Schweisguth, 2004). While Neur was detected in all CCs, we observed a higher level of Neur in the A-CC. Thus, prior to CC intercalation, a Neur-mediated Dl signal produced by the A-CC might activate N in the P-, Pl- and Eq-CCs. Once CCs have intercalated, low N levels were detected apically in CCs, whereas the Dl and Neur patterns remain unchanged. Therefore, following intercalation, the A-CC likely signals to the Pl- and Eq-CCs. To test this, we monitored N signaling by looking at NICD levels over time. To do so, we measured nuclear GFP in the CCs of living NiGFP pupae (Figure 5D and 5E) (Couturier et al., 2012). NICD was detected in the Pl- and Eq-CC before and after CC intercalation. In contrast, NICD was detected in the P-CC only before CC intercalation, suggesting that loss of direct contact with the A-CC results in loss of N receptor activation. Finally, NICD was not detected in the A-CC, further suggesting that this cell is the signal-sending cell within the CC quartet. This pattern of N activity was confirmed by examining the expression of two direct N targets, *E(spl)mδ-HLH* (Figure 5F) and *E(spl)m3-HLH* (not shown), using GFP-tagged reporters. Altogether, our analysis revealed that directional signaling occurs within the CC quartet and that a specific change in the pattern of N activity correlates with CC intercalation.

**Figure 5:**
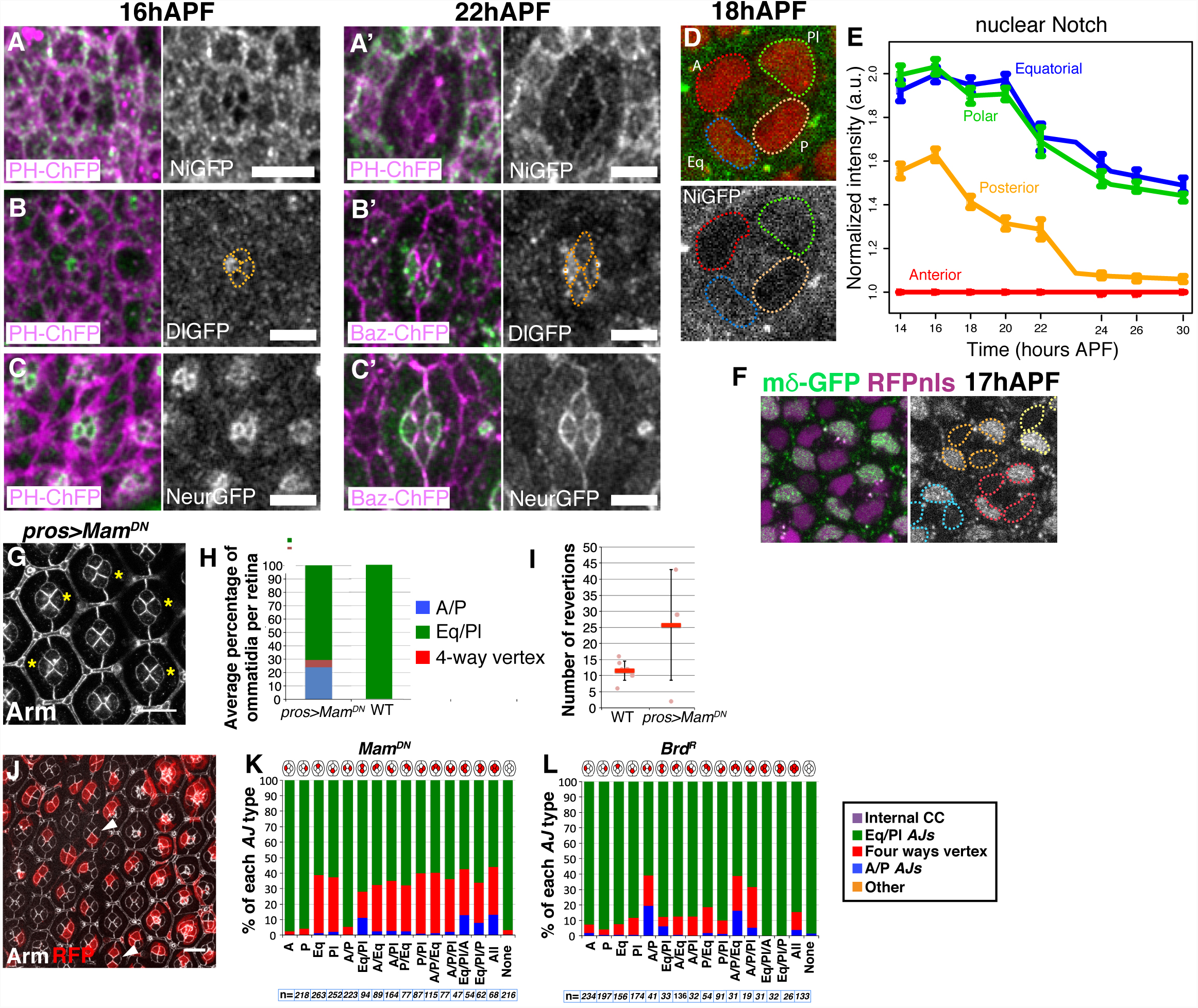
Notch signaling regulates CC intercalation. **(A-A’)** Time-course of NiGFP expression during (Grey) CC intercalation. Cell membranes are labeled with PH-ChFP (Purple). (**B-B’**) Time-course of Dl::GFP expression (grey) during CC intercalation. The CC are outlined using a dashed, orange line. Cell membranes are labeled with PH-ChFP (Purple) in (**B**) and with Baz-ChFP in (**B’**). (**C-C’**) Time-course of Neur::GFP expression (grey) during CC intercalation. Cell membranes are labeled with Baz-ChFP (Purple). (**D**) Representative NiGFP signal (grey) in the CC nuclei, also labeled using an RFP-nls reporter (red). (**E**) Quantification of the nuclear signal for N in the CC cells over time. Note the drop in N signal in the posterior CC as intercalation takes place. (**F**) Representative staining of the N target gene m*δ*-GFP (Grey). CC nuclei are labeled using an RFP-nls reporter (purple). CC nuclei are circled using colored dashed lines with one specific color attributed per quartet. **(G)** *UAS-Mam*^*DN*^ expressed under control of *pros-Gal4*. **(H)** Progression of CC intercalation for retinae expressing *UAS-Mam*^*DN*^ under control of *pros-Gal4* (n=5 retinae, 3433 ommatidia) and for control wild-type flies raised at 25° (n=3 retinae, 1909 ommatidia). **(I)** Number of switches between the different stages of CC intercalation in WT compared to *UAS-Mam*^*DN*^*/pros-Gal4* retinae (WT n=8 ommatidia, pros>Mam^DN^ n=4 ommatidia). **(J)** Single cells expressing *UAS-Mam*^*DN*^ marked by presence of RFP. Arrowheads indicate examples of CCs at a four-way vertex stage when the Eq CC is affected and CCs with a A/P *AJ* when the Eq and Pl CCs are affected. **(K-L)** Quantification of the percentage of ommatidia with each *AJ* type (A/P, four-way vertex, Eq/Pl) when different combinations of CCs express (**K)** *UAS-Mam*^*DN*^, (**L)** *UAS-Brd*^*R*^. N numbers are shown with each panel. Scale bars: **(A-C)** = 5μm, **(G,J)** = 10μm. Error bars: **(I)** = S.D.

To test whether N-signaling contributes to CC intercalation, we used the *UAS-Mam*^*DN*^ transgene, which prevents transcription downstream of N (Giraldez et al., 2002). Expressing Mam^DN^ in all four CCs led to defects in CC intercalation (Figure 5G-H). Live imaging of these retinae revealed that defects in intercalation were due to failures in stabilizing the new Pl/Eq-CC *AJ*, leading to reversion to the original CC configuration (Movie S3 and Figure I). In good agreement with our finding that N is active in the Eq- and Pl-CC, examination of ommatidia mosaic for *Mam*^*DN*^ revealed a requirement for transcriptional regulation downstream of N in the Pl- and Eq-CC in 40% of the cases (Figure 5J-K). To complement this analysis, we also interfered with *Dl* expression by over-expressing a stabilized version of the Neur antagonist Bearded (Brd) (Bardin and Schweisguth, 2006), called Brd^R^ (Perez-Mockus et al., 2017b). Compatible with activation of N in the Pl- and Eq-CC regulating intercalation, we found that expressing Brd^R^ in the A- and P-CCs led to intercalation defects (Figure 5L). These results, together with the finding that Rst is required in the Pl- and Eq-CC during CC intercalation raised the possibility that N signaling promotes CC intercalation by regulating preferential adhesion within the CC quartet and between these cells and the surrounding PPCs.

### Modeling adhesion and MyoII contributions during CC intercalation

To assess the roles of Rst and Hbs mediated adhesion and MyoII driven contractility within the ommatidium, during CC intercalation, we set up and ran vertex model simulations of the ommatidium. We calculated the tension values as weighted averages of MyoII and Rst/Hbs adhesion contributions, with MyoII being directly proportional and adhesion being inversely proportional to the effective tension of a junction. The tension values are calculated independently for intensity measurements before and after the intercalation (Figure 6A-B), normalized to the average of the non-shrinking, non-elongating adjoining CC *AJ*s. Where available, we corrected for the tension values with laser ablations (Movie S4 and S5 and Figure 6C) and applied these tension values on ommatidia clusters of the corresponding topology (Figure 6D for before intercalation topology). We then searched for the tension values that will drive the intercalation in pre-intercalation topologies, and maintain the intercalation in post-intercalation topologies. The tension ranges are obtained from variation of the MyoII averaging weight by 0, 30, 50, 70 or 100 per cent. In addition to varying the junctional tension, we simulated a range of cytosolic MyoII contractility levels corresponding to the PPC, whereby contractility in these cells applies tensile forces on the CC cluster (Figure 6D, dashed red lines; Figure 6E, F, inner myosin mesh).

**Figure 6:**
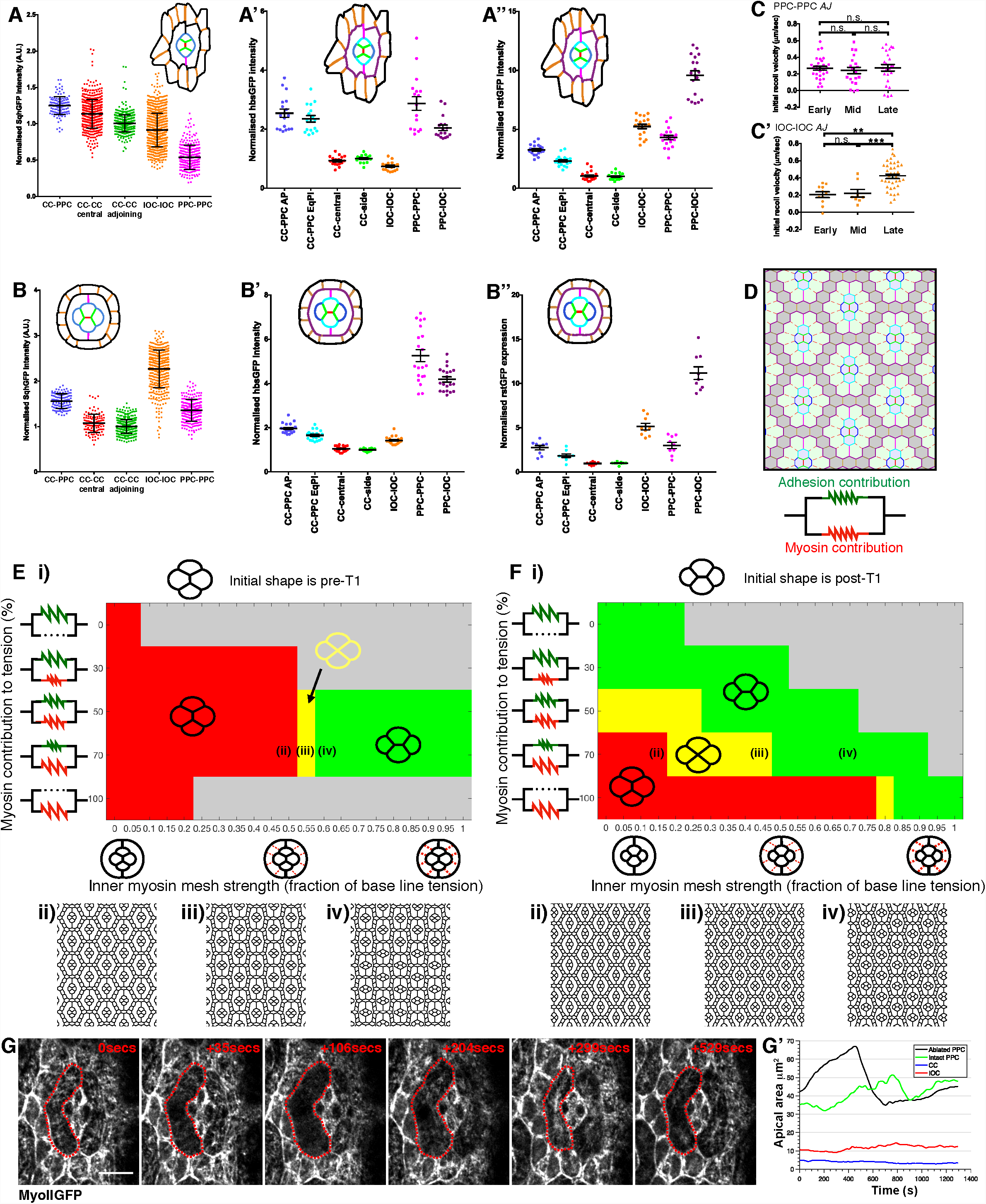
Modeling adhesion and MyoII contributions during CC intercalation. **(A-A’’)** Quantification at *AJ* shrinkage stage of **(A)** Sqh::GFP, **(A’)** Hbs::GFP **(A’’)** Rst::GFP intensity on each *AJ* type normalized to the average of the CC-CC adjoining *AJ* (shown in green). **(B-B’’)** Quantification at *AJ* elongation stage of **(B)** Sqh::GFP, **(B’)** Hbs::GFP **(B’’)** Rst::GFP intensity on each *AJ* type normalized to the average of the CC-CC adjoining *AJ* (shown in green). **(C-C’)** Initial recoil velocity of ablation of **(C)** PPC-PPC *AJs* and **(C’)** IOC-IOC *AJs* at each stage of ommatidial development. For PPC-PPC *AJs*: one-way ANOVA n.s. p=0.784, n=29, 20, 24 *AJs* respectively. For IOC-IOC *AJs*: one-way ANOVA p<0.0001, Tukey’s post hoc: Early-Mid n.s. p=0.97, Early-Late p=0.0001, Mid-Late p=0.0018. n=11, 8, 38 *AJs* respectively. **(D)** The initial geometry for a simulation starts at pre-intercalation topology. Each cell-cell boundary is color-coded following the code used in (**A-C**). The bonds representing the cytosolic contractile actomyosin meshworks are represented as dashed red lines, with the CCs highlighted in blue, PPCs in green and IOCs in grey. The parallel spring schematic represents the tension structure for adhesion and myosin contribution in each cell-cell contact. **(E-Fi)** Simulations started from pre-intercalation **(Ei)** and post-intercalation geometry **(Fi)**. Heatmaps demonstrating the state of intercalation as a function of cytosolic contractile actomyosin meshwork strength, as a function of base tension level (x-axis) and the range of contributions from adhesion and myosin intensity measurements (y-axis). Schematics on top represent the initial geometry at the beginning of the simulation. Spring schematics represent the weight of each adhesion and MyoII in calculation of tension values for each row. Ommatidia schematics represent the strength of the cytosolic mesh. See Methods and Supplementary Table 1 for details. Green represents stable intercalation, red represents failed intercalation (E) or reversion to the initial geometry (F). Yellow represents a stable 4-way junction forming a rosette. Grey points have unstable ommatidia geometry. **(E-Fii)** Simulation snapshots where the tension values cannot drive or stabilize the intercalation. **(E-Fiii)** Simulation snapshots from a 4-way junction forming the rosette. **(E-F iv)** Simulation snapshots from a stable intercalation. The parameters sets for (ii-iv) are marked on heatmaps of (**E-F**). (**G**) Laser ablation of the cytosolic contractile actomysoin meshworks in the PPCs. All cells express Sqh::GFP to visualize MyoII. The targeted cell is outlined in red. (**H**) Measurements of the apical area of the ommatidial cells upon ablation and repair showing that the targeted PPC is under tension. Error bars: **(A-B)** = S.D., **(C)** = S.E.M.

Our simulations suggest that tensions based on solely MyoII or solely on Rst /Hbs adhesion levels cannot drive CC intercalation (Figure 6E, top and bottom rows). Starting from the pre-intercalation topology, CC intercalation can only be simulated when the contributions of adhesion protein and MyoII levels are considered together (Figure 6E, rows for MyoII contribution 50 and 70 per cent). The stability of the topology after intercalation has a larger stable range, in that there are more possible stable set-ups, however the contribution of both MyoII and adhesion must still be taken into account (Figure 6F). In both sets of simulations, the loss of stability in the unstable simulations (grey areas in Figure 6E-F) stems from delamination or intercalation of the IOCs.

One key prediction of the vertex model simulations is that CC intercalation requires some level of tensile force applied by the PPCs to the CC quartet, as shown by the necessity of the inner myosin mesh (Figure 6E-F). This suggests that contractile forces in the PPCs provide pulling forces that contribute in driving stable CC intercalation. To test this prediction, we used a high-power IR laser to perturb the contractile cytosolic meshworks that are present in the PPC (Figure G). Upon ablation, we observed a rapid change of PPC geometry and recoil, followed by a fast repair of the cytosolic meshworks (Figure 6G-G’). These ablation experiments indicate that PPCs are contractile, and are consistent with the prediction that contractile forces outside the CC cluster are required to regulate their intercalation.

## DISCUSSION

The function of most organs depends on the assembly of defined physiological units, which bring together different cell types. Within a physiological unit, the stoichiometry and spatial relationship between cells is usually regulated and underpins the function of the unit. In this context, there are two prevalent modes of epithelial cell movement *in vivo*: intercalation and slithering. Intercalation requires *AJ* remodeling and allows for cells to acquire their niche by moving relative to one another in the plane of the epithelium (Walck-Shannon and Hardin, 2014). Slithering involves epithelial to mesenchymal transition and migration of a cell on top of its neighbors, as for example in the case of single neuroendocrine epithelial cells in the lung epithelium (Kuo and Krasnow, 2015; Noguchi et al., 2015). Here we used the fly ommatidium as a model physiological unit to study a step of developmentally regulated intercalation between four cells. Our work focused on the intercalation of the four glial-like CCs, which occurs over approximately 10 hours. Most instances of epithelial intercalation studied so far have focused on rapid neighbor exchanges, occurring over periods of tens of minutes, in simple epithelia. We found that unlike during rapid cell intercalation, which is powered by the motor protein MyoII, slow CC intercalation is regulated by endocytosis and requires the Rst-Hbs heterophilic adhesion proteins. Our results show that Rst and Hbs are required in the CC quartet for their intercalation. In addition, we discovered that a N-Dl code exists between the four CCs, whereby in the Eq- and Pl-CC, N-signaling is required for CC intercalation to proceed irreversibly. Thus, our work indicates that a N (Eq/Pl-CC) –Dl (A/P-CC) code and an adhesion code through Rst and Hbs regulates CC intercalation. Rst is required in the same CCs as N is, raising the possibility that Rst mediates part of N function during intercalation. Expressing Rst (or Hbs) in Eq/Pl-CC expressing Mam^DN^, does not rescue the defects in CC intercalation (not shown). We therefore envisage that N might regulate the activity of Rst by regulating the expression of a gene that encodes for a regulator of this adhesion protein in the Eq/Pl-CC. In addition, we developed a new vertex model of the ommatidium to study CC intercalation. This model predicted that contractile forces outside of the CCs might be required for these cells to intercalate. Consistent with this prediction, our work indicates that the PPCs are under tension and therefore can exert forces on the neighboring CCs. Altogether, our results show that CC intercalation relies on the interplay of Rst-Hbs mediated adhesion within CC quartet and between the CC and PPCs, and contractile forces in the PPC.

### CC intercalation depends on endocytosis but Myo-II is not essential

Cell intercalation is a main mechanism of epithelial tissue morphogenesis and is typically regulated by the RhoA-Rok-Myo-II pathway, which functions in the corresponding four intercalating cells (Mason et al., 2016; Munjal et al., 2015; Nishimura et al., 2012; Priya et al., 2015; Robertson et al., 2012; Simoes Sde et al., 2010; Simoes Sde et al., 2014). In particular, polarized steps of intercalation that take place over tens of minutes can drive tissue elongation, as seen in the germ-band in *Drosophila* for example. However, our work, which builds upon previous reports that in the fly retina glial-like cells undergo intercalation over approximately 10 hours (Cagan and Ready, 1989; Grillo-Hill and Wolff, 2009), shows that intercalation between these cells presents several departure points from faster modes of intercalation. Firstly, it does not lead to tissue elongation. In fact, the quartet of CCs elongates along the Pl/Eq axis that runs parallel to the shrinking A/P-CC *AJ*, suggesting mechanisms operate that constrain the CC area as intercalation proceeds. In addition, A/P-CC *AJ* shrinkage correlates with an equivalent increase in the length of adjoined Eq/Pl-CC *AJs*, indicating the relative balance amongst the CC-CC *AJs* is actively regulated. Such a compensatory mechanism concomitant to *AJ* shrinkage is not seen in the germ-band for example, where *AJ* shrinkage leads to a decrease in *AJ* perimeter (Bertet et al., 2004). A good candidate for coupling A/P-CC *AJ* shrinkage and lengthening of the adjoined CC *AJs* is the endocytic recycling pathway. As is the case in the germ-band (Levayer et al., 2011), we find that endocytosis is required for CC intercalation, which is consistent with endocytosis being required for shedding membrane during A/P-CC *AJ* shrinkage. Secondly, we find that the Rok, Myo-II, Baz core pathway is largely dispensable for CC intercalation. Similarly, intercalation of tracheal branch cells and squamous epithelial cells in the notum do not require Myo-II (Curran et al., 2017; Ochoa-Espinosa et al., 2017). Instead, in the CCs we find that adhesion through Rst and Hbs regulates this process.

### Neph and Nephrin-like regulates intercalation

Within the developing ommatidium, Rst and Hbs interact in *trans* to regulate the shape of the PPCs and IOCs by favoring adhesion between these two cell types (Bao and Cagan, 2005; Bao et al., 2010). Hbs is required in all four CCs to regulate their intercalation and we could detect low levels of Hbs::GFP expression at the CC/PPC *AJs*. Rst is also detected at these *AJs*, and our genetic experiments indicate that it is required in the PPCs for CC intercalation. Therefore, we propose that at the CC/PPC *AJs*, Hbs comes from the CCs and part of the Rst staining comes from the PPCs. In this model Hbs and Rst are planar polarized in the PPCs, with high levels of Hbs at the PPC/IOC *AJs* and relatively low levels of Rst at the PPC/CC *AJs*. We also note that most of the Hbs-Rst signal is detected at the A-CC/PPC and P-CC/PPC *AJs.* In the case of the Pl/Eq-CCs, only approximately half of the length of their *AJ* with the surrounding PPCs contains Hbs-Rst, suggesting that there is a regulation in these cells that limits the domain of Hbs expression. Altogether, we propose that Hbs-Rst adhesion at the A-CC/PPC and P-CC/PPC *AJs* contributes in occluding the Eq/Pl-CC junction along the Pl/Eq axis of the ommatidium by favoring the A-CC/PPC and P-CC/PPC contacts. This model would also explain why overall the CC quartet lengthens along the Eq-Pl axis of the ommatidium, even though the C/P-CC *AJ* shrinks along this axis.

In addition, while Rst expression and function in the CCs has not been previously reported, our genetic experiments indicate that this protein is required in the Eq/Pl-CCs to regulate CC intercalation. It is difficult to account for this requirement based on the Rst-Hbs preferential adhesion code between the CCs and PPCs. It is however possible that Rst in the Eq/Pl-CCs interacts *in trans* with Hbs presented by the surrounding PPC. We detect only trace levels of Rst and Hbs at the CC-CC *AJs* and whether these trace amounts are sufficient to regulate adhesion between these cells in not clear. It is also notable that Hbs and Rst can interact *in cis* to repress each other’s function (Linneweber et al., 2015). In the case of the Eq and Pl-CCs this might mean that Hbs activity is lowered when compared to the A- and P-CC where we find no requirement for Rst.

### Notch signaling regulates CC intercalation

We find that within the CC quartet, the CC-specific requirement of Rst in the Eq/Pl-CCs correlates with a requirement for transcription downstream of the N receptor. N-signaling has previously been shown to regulate the pattern of Rst/Hbs expression between the PPCs and IOCs in the fly retina (Bao, 2014). N receptor activation is required in the PPCs for Hbs expression, and ectopic expression of N in the IOCs is sufficient to induce Hbs expression and repress Rst in these cells (Bao, 2014). However, it is at present not clear whether Rst and Hbs are direct targets of the N-signaling pathway. Altogether, our results raise the possibility that N acts at least in part through the preferential adhesion system in the CC. Our genetic experiments confirm that Hbs is required in all 4 CC and therefore, it is unlikely that during CC intercalation N instructs Hbs expression. It will be interesting to decipher how exactly N-signaling regulates intercalation in this model system

### Forces external to the intercalating quartet are required for intercalation

MyoII is a key surrogate for tensile strength in cell-cell junctions. Here our vertex model demonstrates the physical balance of forces for stable CC intercalation can only be captured when both Hbs/Rst mediated adhesion and MyoII intensities are jointly taken into account. We also find that the neighboring PPCs are required to provide additional forces to drive neighbor exchange. Our simulations and laser ablations are consistent with contractility in the PPCs regulating CC intercalation through promoting pulling forces onto the A/P-CC.

## Supporting information

Mov S1

Mov S2

Mov S3

Mov S4

Mov s5

## Acknowledgements

We are grateful to Tiffany Cook for providing us with the *prosGal4* line and plasmid, and to the Pichaud lab and Shiladitya Banerjee for discussions related to this study. Stocks obtained from the Bloomington Drosophila Stock Center (NIH P40OD018537) and the Vienna Drosophila Resource Center were used in this study. This work was funded by an MRC grant to F. Pichaud (MC_UU_12018/3). M. Tozluoglu is funded by a Sir Henry Wellcome Fellowship (Grant No: 103095). Y. Mao is funded by an MRC Fellowship MR/L009056/1, a UCL Excellence Fellowship, a NSFC International Young Scientist Fellowship 31650110472 and a Lister Institute Research Prize Fellowship. The work of F. Schweisguth’s lab is supported by FRM-DEQ20180339219 and ANR-10-LABX-0073 grants. M. Trylinski is supported by an ARC fellowship.

## MATERIALS AND METHODS

### Fly strains

Flies were raised on standard food at 18°C. Crosses were performed at 25°C or 29°C as stated. The following fly strains were used:

*;Ecad::GFP;* (Huang et al., 2009).

*sqh*^*AX*3^*; sqh>sqh::GFP;* (BL #57144, (Royou et al., 2002)).

*;Sp/CyO;pros-Gal4/TM6* (Gift from T. Cook).

*;GMR-Gal4*; (Freeman, 1996).

*;;UAS-shibire*^*ts1*^ (BL #44222, (Koenig and Ikeda, 1983)).

*;UAS-YFP-MyoII*^*DN*^; (Barros et al., 2003).

*y,w,rok*^*^2^*^, *FRT19A/FM7;;* (BL #6666).

*y,w,baz*^*xi*^, *FRT9.2/* (Nusslein-Volhard et al., 1987).

*;FRT40A, Ncad*^*M*19^*/CyO; (Iwai et al., 1997)*.

*ubiGFP,FRT19A;eyFlp.*

;*ubiGFP,FRT9.2;eyFlp.*

;*eyFlp;FRT40,GMR-myrRFP/CyO; rst::GFP;;* (BL #59410).

*;hbs::GFP;* (BL #65321).

*;UAS-hbsRNAi;* (VDRC #105913, VDRC #40896).

*;UAS-rstRNAi;* (VDRC #951, VDRC #27223).

*hsflp;;act>CD2>GAL4,UAS-RFP* (BL #30558).

*hs-flp;actin>y>gal4,UAS-mCherry;armGFP/TM6*.

*;;UAS-Mam*^*DN*^ (BL #26672, (Helms et al., 1999).

*w;;UAS-brdKtoR* (Perez-Mockus et al., 2017b).

*Neur::GFP*, a BAC transgenic line with two copies of GFP-tagged Neur (Perez-Mockus et al., 2017b).

*ubi-Baz::mCherry* (Bosveld et al., 2012);

*UAS-Brd*^*R*^ (Perez-Mockus et al., 2017b).

*Dl::GFP*, a GFP knock-in allele (Corson et al., 2017).

*Ni::GFP*, a GFP knock-in allele (Trylinski et al., 2017).

*E(spl)m3-HLH--GFP*, a GFP knock-in line (Couturier et al., unpublished).

*E(spl)mδ-HLH*-*GFP*, a BAC transgenic line expressing GFP-tagged E(spl)m *δ* -HLH (Couturier et al., unpublished).

*w;;FRT82B, ubi-nlsRFP* (BL-30555)

### Clonal analysis

To generate single cell clones, *hs-flp;;actin>CD2>gal4,UAS-RFP* or *hs-flp;actin>y>gal4,UAS-mCherry;arm-GFP/TM6* were crossed to *UAS-* transgenes of interest. Flies were heat shocked at third instar larval stage at 37°C for 10-15 mins and then dissected 2-3 days later at 40%APF (25°C).

### Inhibition of endocytosis using shibire^ts1^

*;;UAS-shibire*^*ts1*^ flies were crossed to *prosGal4* flies and raised at 25°C until the stated time of development, then transferred to 31°C to block endocytosis for either 4hrs or overnight. Retinae were dissected at 40% APF and scored for progression of CC intercalation.

### Antibodies and immunostaining

Pupae were staged at 25°C or 29°C to 40%APF then retinae were dissected in PBS on ice and fixed in 4% paraformaldehyde for 20mins at room temperature (RT). Retinae were washed in PBS-Triton 0.3% (PBS-T) then stained with primary antibody in PBS-T for 2hrs at RT or overnight at 4°C. Retinae were washed in PBS-T and then stained with secondary antibodies for 2hrs at RT. Retinae were mounted in Vectashield (Vectorlabs). The following antibodies were used: mouse N2 A71 anti-Armadillo (1:50), deposited to the DSHB by Wieschaus, E. (DSHB Hybridoma Product N2 7A1 Armadillo) (Peifer and Wieschaus, 1990) and DCAD2 anti-E-Cadherin (1:50), deposited to the DSHB by Uemura, T. (DSHB Hybridoma Product DCAD2) (Yoshida-Noro et al., 1984) combined with mouse or rat secondary antibodies conjugated to Dy405, Alexa488, Cy3 or Alexa647 (Jackson ImmunoResearch) as appropriate, used at 1:200. Images of fixed retinae were acquired on a Leica SPE, Leica SP5 or Leica TCS SP8 confocal microscope. A 40x oil objective was used for imaging of whole retina for quantification and a 63x oil objective was used for higher magnification images.

### Image processing

All images presented were processed using FIJI (Schindelin et al., 2012) and AdobePhotoshop CS4 (Adobe). Graphs were produced in Excel (Microsoft), GraphPadPrism 7 (GraphPad) or MATLAB R2017a (Mathworks). Figures were mounted in Adobe Illustrator CS4 (Adobe).

### Statistical tests

Statistical tests were performed in GraphPad Prism 7. Data was compared using a T-test (paired or unpaired as appropriate) or One-way ANOVA with Tukey’s post-hoc tests.

### Time-lapse imaging

*;Ecad::GFP;, ;Ecad::GFP/Ecad::GFP;UAS-MamDN/pros-Gal4 or ;Ecad::GFP/+;UAS-shibire^ts1^/pros-Gal4* flies were staged to between 10-20%APF at 25°C and the pupal case removed at the dorsal end to expose the retina. Pupae were mounted on blue-tac with the retina facing upwards and covered with a coverslip as previously described (Fichelson et al., 2012) and (Couturier et al., 2014). Time-lapse imaging was performed on a Zeiss inverted microscope with an Andor spinning disc using a Plan Neofluar 100x/1.3 Ph3 oil immersion objective. Images were acquired using ImageJ Micromanager software (Edelstein et al., 2014). Retinae were imaged for a minimum of 12hrs taking a Z-series in 1μm sections every 5mins. Drift in XY and Z was corrected manually. Images were post processed in FIJI to further correct for drift. For *;Ecad::GFP/+;UAS-shibire*^*ts1*^*/pros-Gal4*, flies were raised at 25°C until 15-20% APF and then transferred to the microscope and incubated at 31°C to stimulate endocytosis inhibition as soon as imaging began. NICD levels were measured over time in *NiGFP* pupae using small z-stacks centered at the nucleus level. Image acquisitions were performed at 20 ± 2°C, using a laser scanning confocal microscope (LSM780; Zeiss) with a 63x (Plan APO, N.A. 1.4 DIC M27) objective.

### Laser ablation

*;Ecad::GFP* pupae were raised and mounted as for time-lapse imaging experiments. PPC-PPC and IOC-IOC *AJ* ablation were performed using a Zeiss LSM880 microscope with a Plan Apochromat 63x/NA1.4 oil objective using 740nm multiphoton excitation from a Ti-sapphire laser. An ROI of 3×3 pixels was drawn over an *AJ* and ablated with 5-10% laser power at the slowest scan speed for 1 iteration. Images were acquired every 1sec after ablation. Settings were optimised by imaging *sqh*^*AX3*^*;sqh::GFP;* flies during ablation to ensure only the *AJ*-associated Myo-II was removed and that the medial meshwork of Myo-II remained intact. The *AJs* repaired after every instance of ablation, indicating that the cell was not damaged. Positions of the two adjoining vertices after ablation were manually tracked using FIJI and the distance between them calculated at each frame after ablation. Recoil velocity was calculated by a linear fit across the first frames after ablation. One-way ANOVA was performed in GraphPad Prism7 to compare stages.

### CC area through time

The Tissue Analyser FIJI plugin (Aigouy et al., 2010) with manual correction was used to segment the CCs on a time-lapse (5mins/frame) of *;Ecad::GFP;* retinae. The areas of individual CCs were then measured and averaged across four ommatidia.

### CC axes length through time

The CC-PPC *AJ* perimeter of the CC cluster was traced manually using the Freehand selection tool in FIJI on every 20th frame (100mins) of time-lapse of *;Ecad::GFP;* retinae. To measure A/P and Eq/Pl axes length an ellipse was fitted over the CCs, measured over time and then expressed as a ratio. Measurements were averaged for each time point over 13 time-registered ommatidia from two independent retinae.

### *AJ* perimeter measurements

*AJ* lengths were taken as the inter-vertex distance and were measured manually on each frame of time-lapse movies of *;Ecad::GFP;* retinae using the line tool in FIJI. Measurements were averaged from 13 time-registered ommatidia from two independent retinae.

### Cross-correlation of *AJ* length

Curves of *AJ* length over time for the CCs were smoothed by taking a five point moving average. Data were detrended by taking the running difference to find the change in *AJ* length over time. Cross-correlation was performed in R statistical package using the ccf function. The mean cross-correlation function was calculated as the average correlation coefficient at each time lag across 13 ommatidia from two independent retinae.

### *AJ* Myo-II intensity

*sqh*^*AX3*^*;sqh-sqh::GFP;* flies were staged to 20% APF or 30% APF and retinae were dissected, fixed and stained against Arm. Retinae were imaged on a confocal microscope (Leica SP5) using the same settings for each retina. A Z-projection was made over the depth of the *AJs* in FIJI. A 7 pixel wide line was drawn over each *AJ* in the Arm channel and the intensity of each channel was averaged over this line, except for the CC-PPC *AJs*, as the Sqh::GFP was found to be present only on one side of the *AJ*, a 4 pixel (∼ half of 7) line was drawn over the Sqh::GFP channel. For 20%, adjoining CC-CC *AJs* intensity was averaged within each ommatidium. To control for variations in staining and/or imaging, all individual values within each image where normalized by dividing by the average value for the adjoining CC-CC *AJs* for that image. One-way ANOVA followed by Tukey’s post hoc tests for pairwise comparison was performed using GraphPad Prism 7. For comparison of CC-CC *AJs*, a paired T-test was performed paired by ommatidia.

### Scoring of progression of CC intercalation

Retinae were dissected at 40% APF, fixed and stained for Arm to label the *AJs*. Whole retinae were imaged with a 40x objective taking a Z-series in 0.5μm slices in two tiles. Tiles were manually aligned and combined using Align3 TP plugin in FIJI. Ommatidia were manually scored for stage of CC intercalation across whole retina. Stage was determined by comparison of the position of the central CC-CC *AJ* and the PPC-PPC *AJs*: parallel = *AJ* shrinkage stage and perpendicular = *AJ* elongation stage. Percentages were calculated per retina and then the average percentages and standard deviations were obtained for each genotype. For large clones, ommatidia were compared from inside and outside the clones. For single cell clones, relative position of cells (A, P, Eq and Pl) was recorded and ommatidia were grouped into categories by which cells were affected.

### Scoring switches between stages of CC intercalation

For the *;Ecad::GFP;* and *;Ecad::GFP/Ecad::GFP;UAS-Mam*^*DN*^*/pros-Gal4* time-lapse movies, ommatidia were scored for which stage of CC intercalation they were in for each frame of the movie (5min intervals). A ‘switch’ in a given frame is defined as being scored as at a different stage to that of the previous frame. The total number of switches was measured from 10% of retinal development (before CC intercalation) for roughly 12hrs (until after CC intercalation in WT).

### Particle Image Velocimetry (PIV)

*sqh*^*AX3*^*;sqh-sqh::GFP;* ommatidia were imaged on a Zeiss LSM880 microscope with a Plan Apochromat 63x/NA1.4 oil objective using airyscan detectors to increase resolution. Movies were processed by Bleach correction, Gaussian blur and registered with the Stack-reg plugin (Thevenaz et al., 1998) if needed in FIJI. PIV analysis was performed using the FIJI PIV plugin (Tseng et al., 2012), by choosing an 8×8 pixel window with a time lag 4.34 seconds. Cell contours were tracked using the Tissue Analyzer plugin (Aigouy and Le Bivic, 2016) in FIJI or manually in FIJI to segment the CCs. The angles of each PIV vector within the CC over time were plotted in a polar histogram in MATLAB to show lack of overall directional flow.

### Theoretical modeling

The ommatidium is modelled using the well-established cellular vertex model (Farhadifar et al., 2007). In vertex models, the cells are described by the polygons formed of their contacting edges. These edges represent the attached contacting surfaces (membrane and actomyosin cortices) of both cells. The positions of vertices forming all edges are traced, and the energy minimization is carried out over these positions. The energy of the system is defined by the combined energy contributions of: i) cell area conservation, the deviation of each cell from its ideal size, ii) combination of tension and adhesion energies at each contacting cell-cell boundary (1).

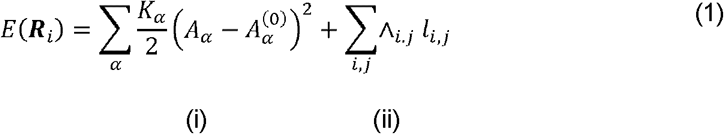

Here, ***E***(***R***_***i***_) is the total energy of the system for a given set of vertex position (***R***_***i***_) that the algorithm minimizes. The system is composed of *N*_*c*_ cells (a = 1…*N*_*c*_), and *N*_*v*_ vertices (***i*** = 1…*N*_*v*_). For area conservation term (i), *K*_α_ is the elasticity coefficient, A_α_ is the current area of the cell, and A^(0)^_α_ is the ideal area. For the tensile/adhesive contact energy contribution (ii), Λ_ij_ is the line tension coefficient for the junction couple (i,j), and *l*_***ij***_ is the length of the junction between vertices. For the simulations in this manuscript, the base tension is Λ_ij_ = 0.26, base A^(0)^_α_ is 1.0 (see Appendix-Table 2 & Fig. EV2B), and *K*_α_ is 1.0 and *K*_α_ is 1.0. The small cone cell area is set to A^(0)^_α_ and remaining cell sizes are scaled accordingly with experimental size scaling measurements (Sup. Table 1). The symmetrical side junctions in between the cone cells are selected as the base, similar to experimental intensity measurements depicted in Figure 6A-B. A weighted average of the adhesion and myosin intensity measurements (Figure 6A&B) are used as surrogate for scaling tension values (Sup Table 1). To scale the tension contribution of adhesion to a cell-cell junction, the base tension level is scaled inversely to normalized adhesion intensity of the junction. For scaling myosin levels, the tension is scaled directly proportional to normalized myosin intensity. This is given in the following equation:

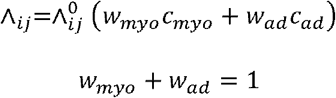

Here, (w) are the averaging weights of myosin and adhesion and (c) are the normalized intensities. For the laser ablated connections, in the pre-intercalation (early) phase, PPC bond, and IOC-IOC bonds on the sides and top & bottom of the ommatidium were not significantly different, therefore were equated to PPC tension value. During the post-intercalation (late) phase, the PPC contact and IOC-IOC contacts on the sides of the ommatidium were not significantly different, whereas the tension on the top & bottom IOC-IOC contacts was 51 percent higher (Figure 6C). The laser ablations depict the true tension of a bond, and model parameters are scaled accordingly where information is available (Sup Table 1). Simulations are carried out on a setup of 91 connected ommatidia with fixed boundaries, and analysis is done on the central ommatidium.

**Supplementary Table 1.**

| Cell type | Ideal area as a fraction of $A_a$ | |
| --- | --- | --- |
|  | Pre-intercalation | Post-intercalation |
| Pigment cell (PPC) | 11.4 | 8.7 |
| Cone cell (CC) sides | 1 | 1 |
| Cone cell (CC) top/bottom | 1.3 | 1.3 |
| (IOCs) | 1 | 1 |

| Contact type | Tension as a fraction of $\Lambda_{ij}^0$ *corrected for laser ablations | | | | | | | |
| --- | --- | --- | --- | --- | --- | --- | --- | --- |
|  | Pre-intercalation |  |  |  | Post- intercalation |  |  |  |
| | $c_{ad} - \text{hbs based}$ | $c_{myo}$ | Sample $w_{ad} = 1$ | Sample $w_{myo} = 1$ | $c_{ad} - \text{hbs based}$ | $c_{myo}$ | Sample $w_{ad} = 1$ | Sample $w_{myo} = 1$ |
| CC-CC side contact | 1.00 | 1.00 | 1.00 | 1.00 | 1.00 | 1.00 | 1.00 | 1.00 |
| CC-CC central contact | 0.93 | 1.10 | 1.07 | 1.10 | 1.05 | 1.06 | 0.96 | 1.06 |
| CC – PPC (EqPI) | 2.35 | 1.30 | 0.43 | 1.30 | 1.65 | 1.52 | 0.60 | 1.52 |
| CC – PPC (AP) | 2.54 | 1.30 | 0.39 | 1.30 | 1.96 | 1.52 | 0.51 | 1.52 |
| PPC-PPC contact | 2.87 | 0.50 | 0.35 | 0.50 | 5.26 | 1.33 | 0.19 | 1.33 |
| PPC - IOC contact | 2.05 | 0.90 | 0.49 | 0.90 | 4.18 | 1.93 | 0.24 | 1.93 |
| IOC-IOC side contact | 0.74 | 0.90 | 0.35* | 0.50* | 1.43 | 2.24 | 0.19* | 1.33* |
| IOC-IOC top/bottom contact | 0.74 | 0.90 | 0.35* | 0.50* | 1.43 | 2.30 | 0.29* | 2.02* |
| all else | 1.00 | 1.00 | 1.00 | 1.00 | 1.00 | 1.00 | 1.00 | 1.00 |

## FIGURE LEGENDS

**Supplementary Figure 1: Fluctuations in PPC-PPC *AJ* and A/P-CC *AJ* length do not correlate**

**(A)** Length of central CC-CC AJ and the PPC-PPC *AJs* of one ommatidium showing negative correlation. Pearson’s correlation coefficient for this example: r=-0.74 for Pl PPC-PPC and r=-0.67 for Eq PPC-PPC. Average correlation coefficient: r=-0.6±0.19 (mean±S.D.) (n=13 ommatidia). **(B)** Average cross-correlation of rate of change in the length of the central CC-CC *AJ* (shown in blue in schematic) with the PPC-PPC *AJs* (shown in red in schematic) (n=13 ommatidia). Error bars = S.D.

**Movie S1: Time-lapse of a representative CC intercalation.** Cells are labeled using ECad::GFP. Frame interval = 5 minutes. Scale bar = 5μm.

**Movie S2: Time-lapse of a representative CC intercalation upon endocytosis inhibition.** Cells are labeled using ECad::GFP. Frame interval = 10 min. Scale bar = 5μm.

**Movie S3: Representative CC quartet expressing Mam^DN^.** Cells are labeled using ECad::GFP. Frame interval = 5 minutes. Scale bar = 5μm.

**Movie S4: Representative laser ablation of an Inter-IOC *AJ***. Cells are labeled using ECad::GFP. Frame interval = 1 second. Scale bar = 5μm.

**Movie S5: Representative laser ablation of an Inter-PPC *AJ***. Cells are labeled using ECad::GFP. Frame interval = 1 second. Scale bar = 5μm.

