## Supplementary figures and images for "Neph/Nephrin-like adhesion and tissue level pulling forces regulate cell intercalation during *Drosophila* retina development"

### Fig S1

#### Supplementary Figure 1: fluctuations in PPC-PPC *AJ* and A/P-CC *AJ* length do not correlate

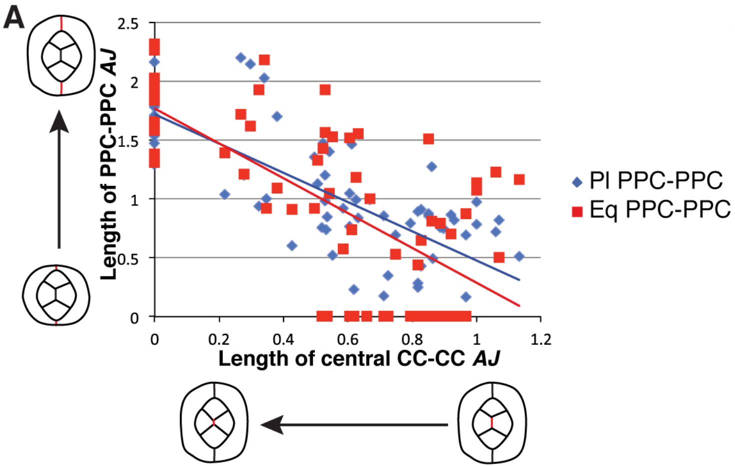


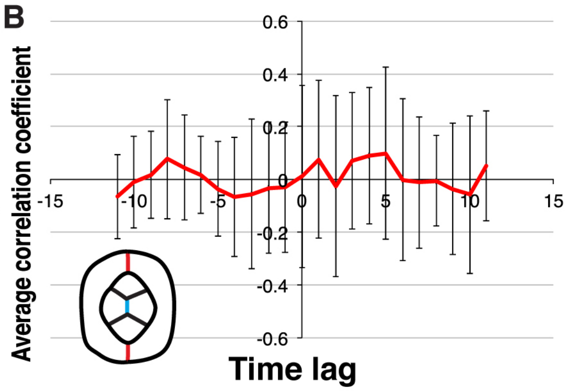
